# Simulated forward and backward self motion, based on realistic parameters, causes strong motion induced blindness

**DOI:** 10.1101/057323

**Authors:** Victoria Thomas, Matthew Davidson, Parisa Zakavi, Naotsugu Tsuchiya, Jeroen van Boxtel

## Abstract

Motion Induced Blindness (MIB) is a well-established visual phenomenon whereby highly salient targets disappear when viewed against a moving background mask. No research has yet explored whether contracting and expanding optic flow can also trigger target disappearance. We explored MIB using mask speeds corresponding to driving at 35, 50, 65 and 80 km/h in simulated forward (expansion) and backward (contraction) motion as well as 2-D radial movement, random, and static mask motion types. Participants (*n* = 18) viewed MIB targets against masks with different movement types, speed, and target locations. To understand the relationship between saccades, pupil response and perceptual disappearance, we ran two additional eye-tracking experiments (*n* = 19). Target disappearance increased significantly with faster mask speeds and upper visual field target presentation. Simulated optic flow and 2-D radial movement caused comparable disappearance, and all moving masks caused significantly more disappearance than a static mask. Saccades could not entirely account for differences between conditions, suggesting that self-motion optic flow does cause MIB in an artificial setting. Pupil analyses implied that MIB disappearance induced by optic flow is not subjectively salient, potentially explaining why MIB is not noticed during driving. Potential implications of MIB for driving safety and Head-Up-Display (HUD) technologies are discussed.

## Introduction

Motion Induced Blindness (MIB) is a visual phenomenon whereby highly salient visual targets become temporarily invisible despite their ongoing physical presence in one’s visual field, when viewed against the background of a global moving mask^1^. MIB is one of many bistable perceptual phenomena^2^ used with increasing popularity to investigate the mechanisms of perceptual organisation^3–5^. These bistable phenomena allow subjects to experience varying phenomenology while receiving physically constant visual input, thus, they can help delineate the neural correlates of consciousness by dissociating physical stimuli from conscious perception^2, 3, 6^.

MIB has been extensively studied under stimulus parameters designed to optimise perceptual disappearances^6–8^. While it has been suggested that MIB may happen in the real world^1^, no research has yet explored MIB using parameters that approximate those experienced in real world movement. To date only two predominantly anecdotal studies have explored whether MIB may occur in the real world. Shimojo^9^ used a mirror ball to create a moving mask of bright spots across a room, and was able to cause the perceptual disappearance of a live person in their interactive museum display. Another demonstration of MIB in real life was reported by Inoue, Yagi and Kikuchi^10^, where they induced MIB by superimposing a target over the optic flow of a movie, filmed from the driver’s point of view while travelling in a car. Perceptual disappearance was induced much more often while travelling in forward motion compared to viewing the target over a still frame. Our own pilot studies have also informally replicated this finding. This evidence suggests that the optical flow experienced in forward self motion can induce MIB in situations where a stable image is projected to the retina over a moving background. In the current study we aimed to lay the groundwork toward establishing the critical parameters of potential MIB phenomena in much more natural situations, focussing on variations of mask speed, trajectory of mask movement and target location.

The most frequently used display for demonstrating MIB consists of three yellow targets and a blue mask of rotating crosses, originally introduced by Bonneh and colleagues^1^. Using this display, various stimulus parameters have been studied^1, 11–16^, among which are the effect of target location^15, 17–21^, mask speed^1, 5, 22^, and cues of mask depth^19, 23^.

In terms of target location, a number of studies have reported greater disappearance when a target is located in the upper left quadrant of the visual display, compared to other tested locations^1, 8, 18–20, 24^. A more frequent disappearance of upper left targets has been replicated several times^1, 17, 19, 22, 24^. However, research finding a significant upper left bias generally uses the original three target triangular display^1^, and finds a significant bias only in relation to the lower central target^1, 18, 24^. There has not yet been a controlled comparison of targets presented in all four quadrants of the screen to determine the effects of visual quadrant on disappearance, despite evidence suggesting that upper visual field may exhibit an effect over target disappearance. The target location effect is an important feature of MIB due to its potential relationship with the neural mechanisms of attention^3, 17^. Indeed, Bonneh and colleagues^1^ interpreted the top-left bias as the result of processing the global mask at the expense of the local target, mediated by the right-hemisphere dominance of visuospatial attention.

It has been shown that mask speed influences MIB, with increased speeds leading to both an increased rate and duration of target disappearance^5, 6^. While it has been suggested that it is the temporal frequency rather than retinal speed of the mask underlying MIB^25^, trajectory of motion has shown to be equally influential^21^. As yet, no study has explored variations in the speed of masks that contract or expand in a way that occurs during self-motion.

Several studies have investigated the effects of other variations in mask composition and movement. In particular, Rosenthal and colleagues^19^ used a three dimensional (3-D) moving mask that mimicked the experience of real world movement and found that significantly more disappearances occurred when masks were perceived to be convex compared to concave, an effect restricted to targets presented in the left field of vision. This and related findings regarding depth ordered masks^23^, suggests that the 3-D interpretation of the visual scene experienced during self-motion is likely to influence MIB, prompting a need for further research on how optic flow (with its implicit depth cues) may influence target disappearances.

We aim therefore to explore whether mask properties modelled on the real world are conducive to MIB, focussing on mask speeds that would typically occur in a driving situation, and the expanding and contracting optic flow experienced in (simulated) forward and backward self-motion. We also aim to explore the effect of target location on disappearance, to investigate the previously identified upper left bias for disappearance and to test the hypothesis that MIB is mediated by the right-hemisphere dominant attentional mechanism.

Finally, while it is possible that MIB might be occurring in real-life settings, it has not yet been reported. A reason for this may be that the saliency of MIB phenomenon is so low that we do not notice it in everyday life. To estimate the saliency of MIB, phasic pupillary responses have recently been investigated during MIB events^26^. It was found that MIB produced larger pupillary responses than target re-appearance, suggesting disappearance is a more subjectively salient and surprising event to the brain^26–29^. Previous research however, has not manipulated mask type to examine if such an interpretation of pupillary response as a proxy of subjective saliency holds across different stimulus conditions. Therefore we will also investigate the saliency of target disappearance and reappearance in our stimuli, measured by phasic pupil response, as well as the overall saliency of the different movement patterns by looking at tonic (i.e., average) pupil dilation for each mask condition.

## Results

### Experiments 1 and 2: MIB can be induced by the optic flow of simulated forward and backward self-motion

We investigated the duration of MIB disappearance with optic flow patterns in two sets of non-overlapping subjects (Experiment 1: *n* = 18; Experiment 2: *n* = 19, see Method) comparing masks simulating forward (expansion) and backward (contraction) self motion with a static stimulus (see Figure 1 and Supplementary Video S1 for experimental display). The design of Experiments 1 and 2 were identical in all aspects, with the exception of the use of an eye-tracking device in Experiment 2 to monitor eye movements (as well as a small variation in viewing distance).

**Figure 1.**
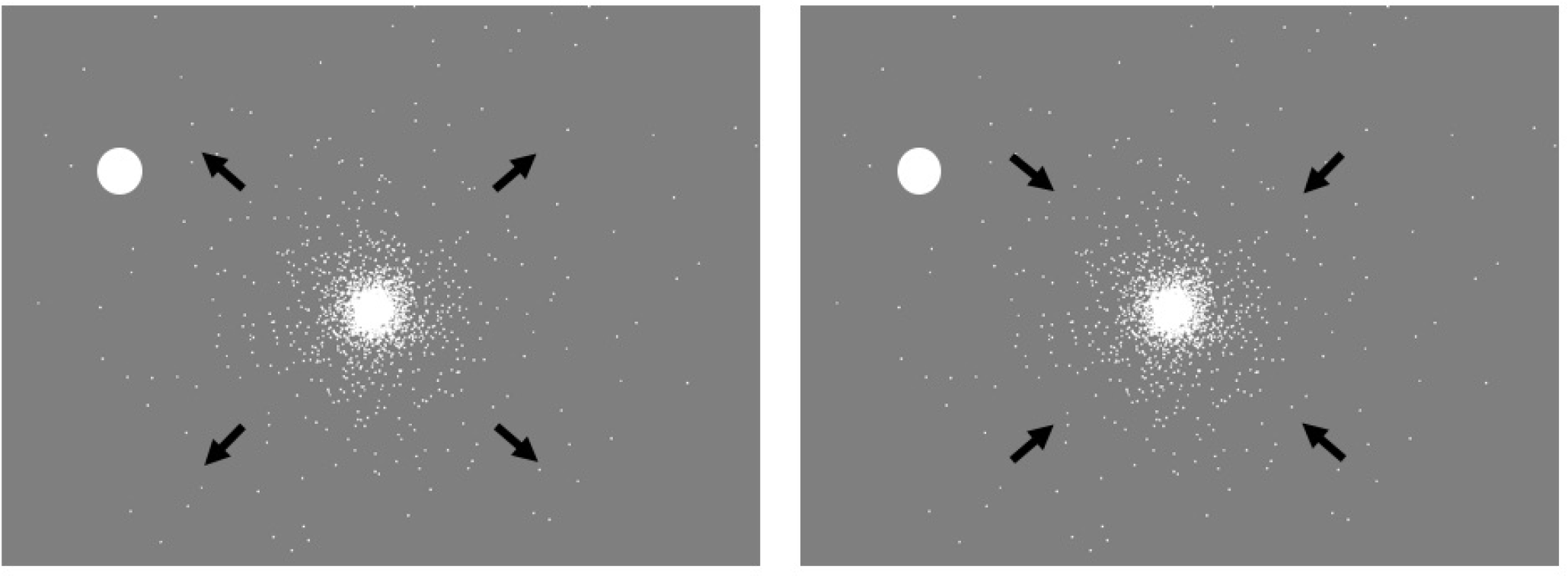
Experimental display for Experiments 1, 2 and 3. Schematic black arrows were not shown on the display. They depict expanding motion shown on the left, and contracting motion shown on the right.

In Experiment 1, repeated measures ANOVAs found a significant main effect of mask type on mean percentage and mean rate of target disappearance (see Table 1). Experiment 2 replicated this finding for both mean percentage and mean rate of target disappearance (see Table 1). As the distribution of perceptual disappearances have been shown to follow a non-normal gamma distribution^26^, we also analysed the effect of mask type on the median percentage and rate of target disappearance, for which similar results were obtained (see Table 1).

**Table 1.** Repeated Measures ANOVA Results for Experiments 1-4, Using Mean and Median

| Experiment | Analysis | Mean | Median |
| --- | --- | --- | --- |
| Experiment 1: |  |  |  |
| *Optic Flow Mask* | Percentage | $F(2, 32) = 23.30, p < .001$ | $F(2, 32) = 12.23, p < .001$ |
| | Rate | $F(2, 32) = 18.50, p < .001$ | $F(2, 32) = 11.82, p < .001$ |
| Experiment 2: |  |  |  |
| *Optic Flow Mask* | Percentage | $F(2, 36) = 33.30, p < .001$ | $F(2, 36) = 21.23, p < .001$ |
| | Rate | $F(2, 36) = 55.53, p < .001$ | $F(2, 36) = 32.06, p < .001$ |
| Experiment 3: |  |  |  |
| *Mask Speed* | Percentage | $F(3, 51) = 4.90, p = .005$ | $F(3, 51) = 2.44, p = .075$ |
| | Rate | $F(3, 51) = 4.26, p = .009$ | $F(3, 51) = 3.96, p = .013$ |
| Experiment 4: |  |  |  |
| *2D Mask Movement* | Percentage | $F(3, 54) = 19.39, p < .001$ | $F(2.047, 36.845) = 12.55, p < .001$ (GG) |
| | Rate | $F(3, 54) = 15.15, p < .001$ | $F(1.983, 35.699) = 11.66, p < .001$ (GG) |

Note: GG indicates Greenhouse-Geisser correction of degrees of freedom due to violated Sphericity

Post-hoc paired-samples t-tests for Experiment 1 (Figure 2; using Bonferroni-corrected p-values) revealed that mean percentage of target disappearance (i.e. the percentage of MIB) in the static control condition was significantly lower than during both expansion (*t*(16) = 5.65, *p <*.001, *r* = .82) and contraction (*t*(16) = 5.45, *p <*.001, *r* = .81). Mean rate of target disappearance was also significantly lower than during both expansion (*t*(16) = 4.53, *p* = .001, *r* = .75) and contraction (Figure 2; *t*(16) = 5.27, *p <*.001, *r* = .80). Post-hoc paired-samples t-tests for Experiment 2 (Figure 3; using Bonferroni-corrected p-values) also found mean percentage of target disappearance in static mask conditions to be significantly lower than during both expansion (*t*(18) = 6.01, *p <*.001, *r* = .82), and contraction (*t*(18) = 6.57, *p <*.001, *r* = .84). Mean rate of target disappearance during a static condition was also significantly lower than during both expansion (*t*(18) = 7.50, *p <*.001, *r* = .87) and contraction (*t*(18) = 9.70, *p <*.001, *r* = .92) (Figure 3). Mean percentage of target disappearance for expansion and contraction were not significantly different from each other in Experiment 1 (*p* = .48) and Experiment 2 (*p* = .42). Likewise, mean rate of disappearance for expansion and contraction were not significantly different from each other in Experiment 1 (*p* = .94) and Experiment 2 (*p* = .87).

**Figure 2.**
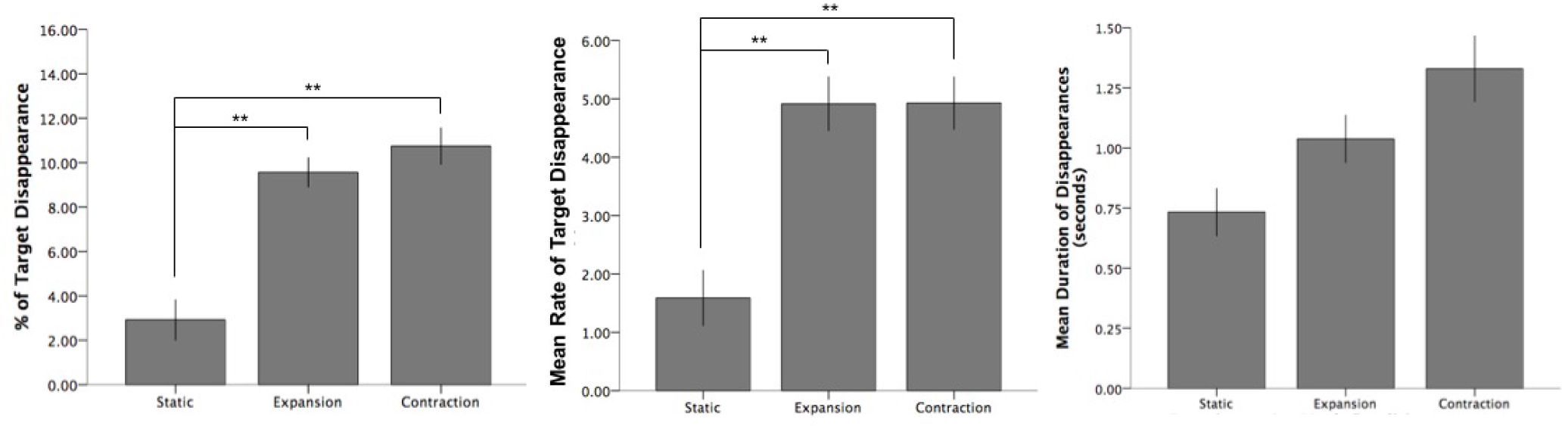
Experiment 1. Percentage, Mean Rate and Mean Duration of target disappearance dependent on movement type. Double asterisk (**) indicates significance at the .01 level; error bars indicate standard error (+/− 1 SE)

**Figure 3.**
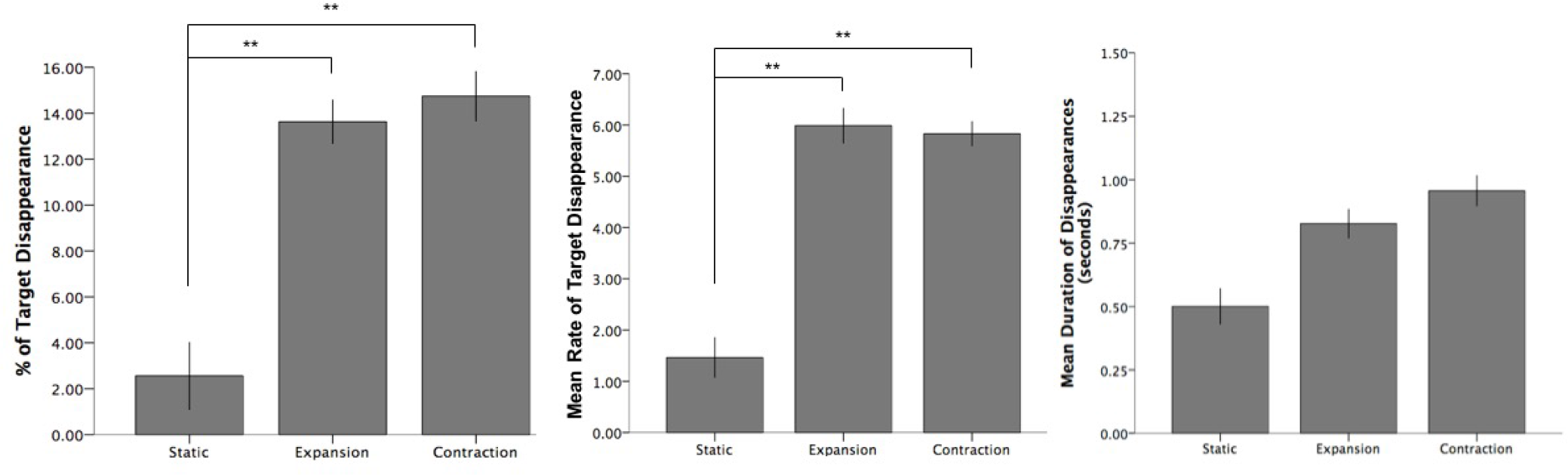
Experiment 2. Percentage, Mean Rate and Mean Duration of target disappearance dependent on movement type. Double asterisk (**) indicates significance at the .01 level; error bars indicate standard error (+/− 1 SE).

In terms of length of individual disappearance episodes, no significant difference was found between conditions on mean (*F*(2, 24) = 3.22, *p* = .06) or median (*F*(2, 24) = .65, *p* = .53) analyses in Experiment 1 (Figure 2), or mean (*F*(2, 24) = 3.31, *p* = .05) or median (*F*(2, 24) = 1.28, *p* = .30) analyses in Experiment 2 (Figure 3).

### Experiment 2: The influence of saccades and target location

We recorded eye movements with an offline velocity-based saccade detection algorithm^30^ in Experiment 2 and 4, to ensure participants were fixating as instructed, as we expected that saccades towards the target would reduce MIB^31^. The direct eye tracking also allowed us to quantify whether our results were attributable to eye movements. For example, it may be possible that the static condition induced more saccades, which then inhibited perceptual disappearances. Our analysis focussed on macro-saccades, because larger saccades have a stronger influence over the visual input, and are most relevant in driving situations. We also confirmed that our results were not due to our selected macro-saccade size (>1°), by also analysing all saccades (micro-saccades + macro-saccades). Figure 4 shows the influence of these saccade effects.

**Figure 4.**
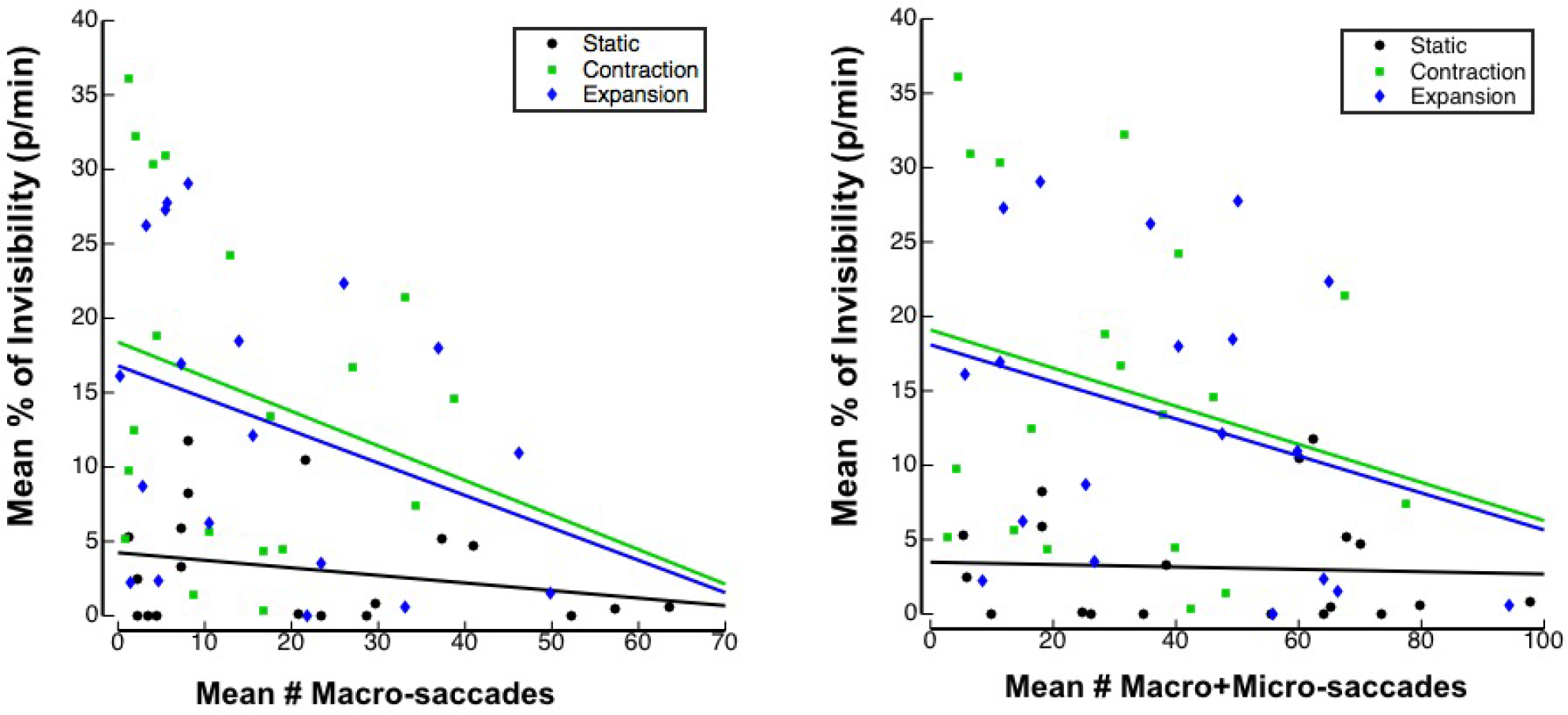
Experiment 2. Relationship between mean number of macro and macro+micro saccades and percentage of disappearance per trial for each mask condition.

Using linear mixed effect (LME) modeling^32^, we quantified how much of the percentage of target disappearance in each trial can be accounted for by the number of saccades, alongside our main factors of interest of mask motion types and target position (see the details of LME in Methods). The significance of each factor was assessed with a likelihood ratio test^32^.

LME and likelihood ratio tests confirmed the significant effects of mask type (Experiment 1: c^2^(1) = 40.71, *p <*.001; Experiment 2: c^2^(1) = 96.92, *p <*.001), even when taking account of the significant effects of saccades (Experiment 2: c^2^(1) = 12.31, *p* = .006). Note the effect size of mask types, measured as c^2^ value^32^, is much larger than the effect of saccades. Thus we conclude that the effects of motion mask types on target disappearance cannot entirely be explained by the saccades in each condition. In addition, we found that the vertical position of the target also influenced the percentage of MIB disappearance (Experiment 1: c^2^(1) = 5.32, *p* = .022; Experiment 2: c^2^(1) = 28.42, *p <*.001) while the horizontal position did not (Experiment 1: c^2^(1) = .95, *p* = .033; Experiment 2: c^2^(1) = .024, *p* = .63). See Table 2 for further LME analyses exploring target location with Mean Rate and Median Duration of target disappearance.

**Table 2.** Linear Mixed Model Results for Experiments 1-4, Using Mean Rate and Median Duration of Target Disappearance

| Experiment and Analysis | Mask Type | Vertical Position | Horizontal Position | Saccades |
| --- | --- | --- | --- | --- |
| **Exp 1: Optic Flow Mask** |  |  |  |  |
| Mean Rate | $c^2(1) = 58.11, p < .001$ | $c^2(1) = 2.45, p = .12$ | $c^2(1) = .00066, p = .98$ | |
| Median Duration | $c^2(1) = .73, p = .69$ | $c^2(1) = 1.22, p = .27$ | $c^2(1) = .0024, p = .96$ | |
| **Exp 2: Optic Flow Mask** |  |  |  |  |
| Mean Rate | $c^2(1) = 114.61, p < .001$ | $c^2(1) = 5.53, p = .02$ | $c^2(1) = 2.21, p = .14$ | $c^2(1) = 43.52, p < .001$ |
| Median Duration | $c^2(1) = 9.8, p = .04$ | $c^2(1) = 7.96, p = .005$ | $c^2(1) = 2.17, p = .14$ | $c^2(1) = 3.18, p = .37$ |
| **Exp 3: Mask Speed** |  |  |  |  |
| Mean Rate | $c^2(1) = 15.24, p = .002$ | $c^2(1) = .61, p = .43$ | $c^2(1) = .61, p = .43$ | |
| Median Duration | $c^2(1) = 6.55, p = .09$ | $c^2(1) = 2.65, p = .10$ | $c^2(1) = 2.24, p = .13$ | |
| **Exp 4: 2D Mask Movement** |  |  |  |  |
| Mean Rate | $c^2(1) = 54.302, p < .001$ | $c^2(1) = 1.75, p = .19$ | $c^2(1) = .010, p = .92$ | $c^2(1) = 4.22, p = .38$ |
| Median Duration | $c^2(1) = 21.37, p = .002$ | $c^2(1) = .74, p = .39$ | $c^2(1) = .56, p = .45$ | $c^2(1) = 8.24, p = .08$ |

While the mean number of saccades negatively correlated with the mean MIB percentage across subjects for expansion and contraction mask conditions, the differences between mask types cannot be explained by the number of saccades. This is further shown by a regression analysis with macro-saccades as the independent variable, which showed that 95% confidence intervals of the y-intercepts of both contraction (*b*0 = 18.39, 95% CI [10.21, 26.58]) and expansion (*b*0 = 16.80, 95% CI [9.67, 23.94]) did not overlap with the 95% confidence interval of static (*b*0 = 4.24, 95% CI [1.48, 7.00]). The results of these regression analyses were not significantly changed when performed after combining both macro and micro saccades (Figure 4).

### Experiment 3: MIB is dependent on the speed of simulated forward self-motion

In Experiment 3, we examined the effects of the simulated speed of forward self-motion, comparing masks that simulated moving at 35 km/h, 50 km/h, 65 km/h and 80 km/h. There was a significant main effect of speed on mean MIB percentage and mean MIB rate of target disappearance (see Table 1, with similar results using median values), as seen in Figure 5. Regarding mean percentage of disappearance, post hoc paired-samples t-tests (using Bonferroni corrected p-values) revealed that disappearance percentage for the lowest speed level of 35 km/h was significantly lower than 65 km/h (*t*(17) = 3.59, *p* = .01, *r* = .66). Mean percentage of 35 km/h was also lower than that of 50 km/h (*t*(17) = 2.69, *p* = .09, *r* = .55) and 80 km/h (*t*(17) = 2.90, *p* = .06, *r* = .58), although differences fell short of significance after the Bonferroni corrections applied for multiple comparisons. Similar effects were found in post-hoc tests for mean rate, where number of target disappearances for the lowest speed level of 35 km/h were significantly less than 80 km/h (*t*(17) = 3.08, *p* = .04, *r* = .60) and 65 km/h (*t*(17) = 3.22, *p* = .03, *r* = .62). Mean rate of 35 km/h was also lower than of 50 km/h (*t*(17) = 2.57, *p* = .12, *r* = .53), although the difference fell short of significance after the Bonferroni corrections. We also found that the mean percentage of target disappearance showed linear dependency on the mask speed (*F*(1, 17) = 8.63, *p* = .009, *r* = .58) but neither quadratic (*p* = .08) nor cubic (*p* = .80) dependency was significant, although mean percentage of target disappearance appears to level off at higher speeds. The same effect was found for mean rate of target disappearance, with a significant linear dependency across conditions (*F*(1, 17) = 10.30, *p* = .005, *r* = .61), but neither quadratic (*p* = .18) nor cubic dependency (*p* = .49).

**Figure 5.**
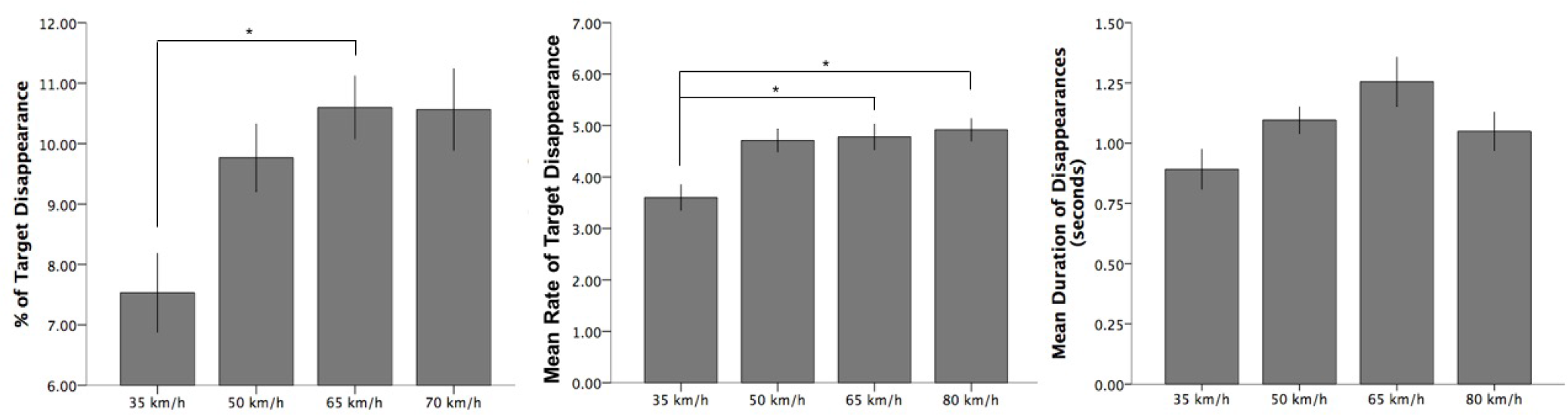
Experiment 3. Percentage, Mean Rate and Mean Duration of target disappearance per trial for each mask speed . Single asterisk (*) indicates significance at the .05 level. Error bars indicate standard error (+/−1 SE)

We again performed a linear-mixed effect analysis, investigating the influence of mask type and target location on mean percentage of target disappearance. The analysis showed that mask type had a significant influence (c^2^(1) = 14.58, *p* = .002), while vertical (c^2^(1) = 3.50, *p* = .061) and horizontal position did not have a significant influence (c^2^(1) = 2.87, *p* = .091). See Table 2 for further LME analyses exploring target location with Mean Rate and Median Duration of target disappearance.

In terms of the duration of individual disappearance episodes within Experiment 3, no significant difference was found between the four mask speed conditions on mean (*F*(3, 36) = 2.26, *p* = .10) or median (*F*(3, 36) = 1.88, *p* = .15) analyses (Figure 5).

### Experiment 4: MIB due to static, random and radial mask motion at a constant dot speed

Experiment 4 had two main purposes: 1) to compare our optic flow conditions to a condition with random motion that lacks any coherent motion structure, which has previously been shown to be important in MIB^19, 23^ and 2) to investigate whether MIB still occurred as strongly when mask dots moved at a constant speed compared to a more natural accelerating speed in our previous experiments.

Again, there was a significant main effect of the type of mask movement on percentage (Figure 6) and mean rate of target disappearance (see Table 1, with similar results using median values). Post-hoc paired-samples t-tests (using Bonferroni corrected p-values) revealed that the static control condition had a significantly lower mean percentage of disappearance than during 2-D random (*t*(18) = 5.46, *p <*.001, *r* = .79), 2-D radial expansion (*t*(18) = 5.10, *p <*.001, *r* = .77) and 2-D radial contraction conditions (*t*(18) = 4.80, *p <*.001, *r* = .75). Likewise, mean rate of target disappearance was significantly lower in the static condition compared to 2-D random (*t*(18) = 4.96, *p* = .001, *r* = .76), 2-D radial expansion (*t*(18) = 4.72, *p* = .001, *r* = .74) and 2-D radial contraction (*t*(18) = 4.95, *p* = .001, *r* = .72). However, in both rate and percentage, 2-D random, 2-D radial expansion and 2-D radial contraction were not significantly different from each other.

**Figure 6.**
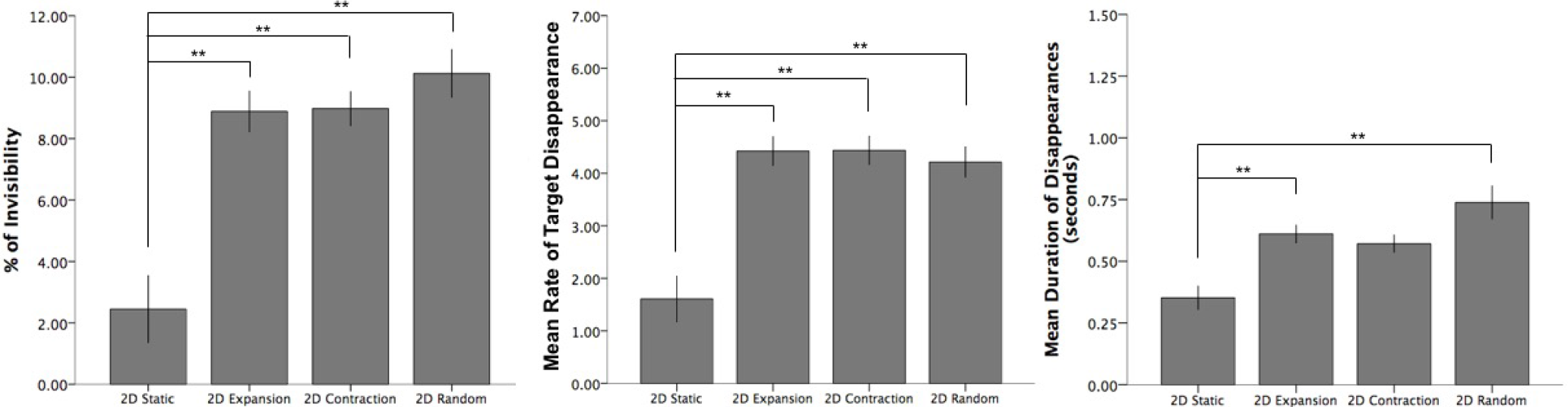
Experiment 4. Percentage, Mean Rate and Mean Duration of target disappearance for each 2D mask movement condition. Double asterisk (**) indicates significance at the .01 level. Error bars indicate standard error (+/−1 SE)

As the mean percentage and rate of disappearance in Experiment 4 appeared to be lower than that of Experiments 1-3, we also compared the data of Experiment 4 to the 35 km/h condition of Experiment 3, which had a mask speed of 3.8°/sec around the target, which closely matched the 3.4°/sec in Experiment 4. This specific comparison between these mask types was necessary because the dot speeds around the location of the target were quite different (i.e., slower near fixation and faster towards periphery) in Experiment 1-3. In this comparison no difference in MIB between conditions was found for either mean percentage (*p* = .50) or mean rate of disappearance (*p* = .50).

Mean length of individual disappearance episodes was significantly different between mask conditions in Experiment 4 (*F*(3, 39) = 7.19, *p* = .001) (Figure 7. Post-hoc paired-samples t-tests (using Bonferroni corrected p-values) revealed that the static control condition had a significantly lower mean length of individual disappearance episodes than during 2-D random (*t*(13) = 3.58, *p* = .003, *r* = .70) and 2-D radial expansion (*t*(13) = 4.12, *p* = .001, *r* = .75). Disappearance length in the 2-D radial contraction condition was also longer than in static (*t*(13) = 2.44, *p* = .030, *r* = .56), although the difference fell short of significance after Bonferroni corrections. For comparison, similar results were found using median length of individual disappearance episodes (*F*(3, 39) = 4.58, *p* = .008).

**Figure 7.**
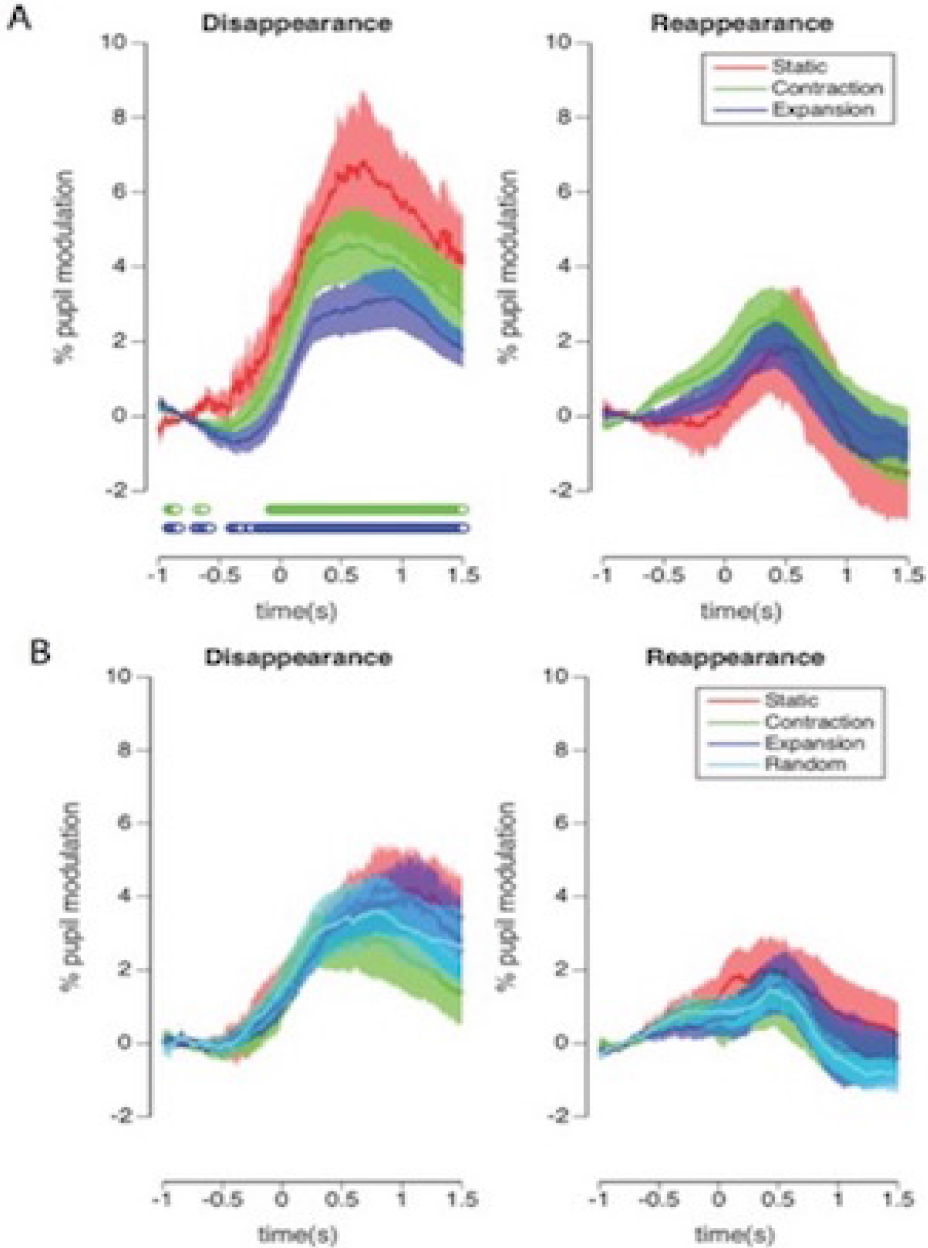
Experiment 2 (A) and 4 (B). Phasic pupil diameter changes (in % modulation) in response to a reported disappearance (left) and reappearance (right). Different conditions are displayed with different colors. Significance is indicated below the plot (Green = contraction is significantly different from static, Blue = expansion is significantly different from static at .05 level. There was no significant difference between contraction and expansion). Shaded areas represent 1 SE.

**Figure 8.**
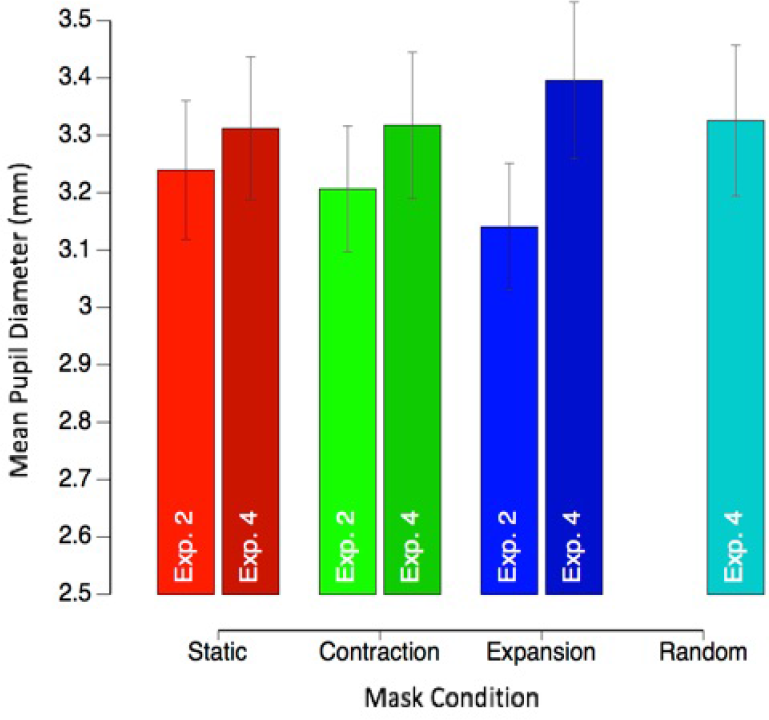
Experiments 2 and 4. Mean tonic (baseline) pupil diameter (+/- 1 SE) per mask condition.

The linear-mixed effect analysis showed that saccades overall did influence percentage of MIB disappearance (c^2^(1) = 9.62, *p* = .047). However the effect size was much smaller than the highly significant influence of mask type (c^2^(1) = 68.85, *p <*.001). Vertical position of the target (c^2^(1) = 8.13, *p* = .004), but not horizontal position (c^2^(1) = 0.005, *p* = .75) accounted partly for the percentage of MIB disappearance. See Table 2 for further LME analyses exploring target location with Mean Rate and Median Duration of target disappearance.

### Experiments 2 and 4: Phasic pupil responses as a proxy for subjective saliency during MIB disappearance and reappearance

Recent research^26^ has shown that phasic pupil responses are larger for target disappearances than reappearances. We sought to investigate whether this pattern of larger responses to disappearances would be replicated for our optic flow movement, and whether there would be any difference in subjective salience (as inferred from the pupil response) across mask motion types.

In terms of phasic pupil responses, target disappearances induced the largest pupil response in the static condition compared to the contraction and expansion conditions in Experiment 2 (multiple comparisons across time are corrected by FDR in Figure 7), while we found no differences in Experiment 4. The pupil size changes were smaller for target reappearances, consistent with Kloosterman et al^26^, and we did not find any significant differences between reappearance conditions in either Experiment 2 or 4. Our results suggest that perceptual disappearances were subjectively more salient than reappearances, in support of Kloosterman et al^26^, regardless of mask types.

To further test if subjective saliency of the different motion types also modulated the overall tonic (i.e. baseline) pupil diameter, we also compared tonic pupil responses across Experiments 2 and 4, and across motion types (Figure 9). Interestingly, the average pupil diameter seems to correlate with the subjective saliency of the different motion types. Unnatural motion types that violate the optic flow of self-motion (e.g., the expansive 2-D motion in Experiment 4, which lacked an increase in dot speeds at greater eccentricities) showed the largest tonic pupil diameter, followed by 2-D contraction and random motion (Experiment 4). Natural motion stimuli (3-D expansion and contraction, Experiment 2), on the other hand, showed a smaller tonic pupil diameter. Quantitatively, average pupil diameter significantly differed across the conditions of the experiments, (*F*(1, 132) = 6.04, *p* = .001). More specifically, in Experiment 4, average pupil diameter remained comparable to the static condition for contraction (*t*(18) = −0.18, *p* = .86, *r* = .04), but increased significantly for the expansion condition (*t*(18) = −2.44, *p* = .03, *r* = -.50). In Experiment 2 tonic pupil diameter was greater in the static condition, compared to both contraction (not significant; *t*(18) = 0.65, *p* = .52, *r* = .15) and expansion conditions (*t*(18) = 3.43, *p* = .003, *r* = .63). There were no significant differences in phasic or tonic responses dependent on target location, for both disappearance and reappearance (data not shown). We present a potential interpretation of these pupil responses within a predictive coding framework below.

**Figure 9.**
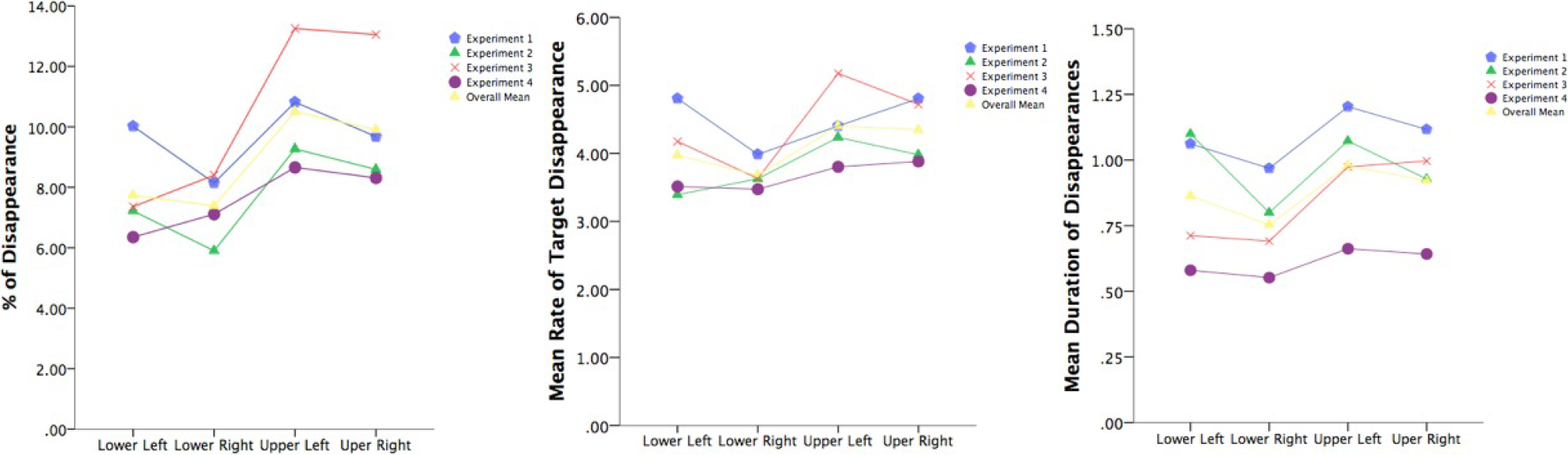
Mean percentage of invisibility per trial according to target quadrant across all experiments. Similar trends were obtained for analyses using mean rate of target disappearance per trial.

## Discussion

In the current study we tested whether stimulus parameters modelled on optic flow simulating self-motion are able to induce MIB. Various other motion patterns have been used in the past^1, 19–21^, but none have used a contracting or expanding optic flow analogous to when walking or driving, despite cues of self-motion being essential to safe navigation^33^. The present study showed that optic flow patterns, similar to those experienced when driving a car in a realistic speed range, can reliably elicit MIB.

### Mask movement

While mask movement was conducive to target disappearance, disappearance was not affected by variations in the direction of movement that we tested within Experiments 1, 2 and 4, with optic flow expansion and contraction causing comparable target disappearance. While this effect may be expected based on the reliability of MIB occurring with a variety of mask parameters^8, 19–21, 23, 25^, it is the first time that optic flow has been shown to elicit MIB. While claims regarding MIB in the real world cannot be made based on this finding alone, it nevertheless functions as a necessary first step in that direction.

Expanding and contracting optic flow were shown to induce significantly more disappearance than radial expansion and contraction patterns with constant speed in Experiment 4, but this difference disappeared once we controlled for mask speed differences between Experiments 1 and 2 vs 4. However, the similar level of disappearance is still surprising, given that the dot density of the mask around the peripheral target within expanding and contracting optic flow was around 9-times lower, compared to 2-D radial expansion and contraction. Lower mask density necessarily results in less competition between the mask and target, causing less target disappearance^1, 20^. Therefore, it is possible that a stronger disappearance effect within simulated forward and backward motion could have been observed if we were to equate dot density around the target in these conditions. Further research addressing inconsistencies between mask movement and dot density around the target will be needed to explore this possibility.

### Mask speed

We found that target disappearance increased with mask speed, consistent with previous research^1, 5^. Specifically, there was a significant positive linear dependency, with upper mask speeds eliciting significantly more disappearances than the lowest of 35km/h in Experiment 3. While overall target disappearance (but not individual episode durations) did increase as a function of mask speed, the differences between the three upper speeds were not significant, suggesting that there may be a plateau effect of increased target disappearance that is reached at around 50 km/h (but, the mean duration of disappearance per episode appears to peak at 65 km/h in Figure 5). Further research is needed with a wider range of speeds to establish whether an optimal range of mask speed for inducing target disappearance exists, and where the upper threshold might be. Such research will extend our current findings regarding mask speed, which holds implications for driving safety, as it is known that driving speed is significantly related to accident rates^34, 35^.

### Target location

To summarise the location biases in MIB, Figure 9 shows the mean duration of disappearance for the different target locations across each experiment. We found significantly greater target disappearance for upper visual field presentation compared to the lower visual field across Experiments 2, 3 and 4 (*p* = .06 for Experiment 1). On the other hand, a bias for the left visual field was not significant in any experiment.

While not significant, there was a trend for target disappearance to be greatest for presentations in the upper left quadrant of the screen which is consistent with previous research^1, 17, 19^. Past research has taken the increased disappearance of upper left targets to suggest a possible neural mechanism underlying MIB^1^, such as spatial attention, which also shows strong visual field asymmetries^36, 37^. However, most studies that report such a bias presented three targets in a triangular display with two targets in the upper field and one centred in the lower field, and found significant results only between the upper left and lower central target^1, 18, 24^. If we interpret this in light of our current findings, in which the upper field bias was much stronger, it is possible that the upper left bias in MIB builds largely on a strong bias to the upper field overall.

The upper field bias of perceptual disappearance is also compatible with previous research which suggests a greater visual acuity exists for the lower visual field^38^, where object recognition in natural scenes is particularly important for self-motion, hunting and general survival^38, 39^. This lower visual field advantage has been reported in a range of visual tasks, including target detection and discrimination^40^, contrast sensitivity^41, 42^, and the perception of illusory contours^43^. The present results extend this list to include MIB, as less target disappearance in the lower visual field may be explained as a consequence of greater visual acuity.

### Saccades and the link to driving

We also considered the possibility that saccades may interfere with MIB. The results of our eye-tracking data showed that saccades influenced target disappearance, consistent with previous research^24, 31^. However, the magnitude of the effect was small compared to the influence of the mask, in both Experiment 2 and 4, and it did not explain our main findings, which is consistent with the observation that saccades do not eliminate MIB entirely^44^ and MIB occurs even with a moving target^1^. This is also evident from our scatter plot analyses (Figure 4), which showed that, at the same level of saccades, the percentage of MIB disappearance was larger in the motion mask types compared to the static masks. Thus, saccades did influence MIB, but cannot explain the difference in MIB between different mask types.

We recorded an average of 35 (macro + micro) saccades per minute in the optic flow mask conditions. Although around 80 saccades per minute typically occur while driving - more than twice as frequent as in our experiments - there is great variation in this according to the type of road and environment^45^. While the frequency of saccades in our experiments were relatively low across subjects, there were some subjects who exhibited 30-40 macro-saccades or 50-70 (macro + micro) saccades per minute (Figure 4) and still reported substantial percentages of target disappearance (10%). As indicated by this percentage, assuming that 5-6 seconds of MIB can occur per minute for the saccade rates reported herein, then up to 10 minutes of MIB may occur during a two hour drive. Further, MIB effects may increase when combined with other effects, such as fatigue and inattentional blindness that are known contributors to accidents^34, 35^. Further studies are needed to address such a possibility in settings which more closely approximate real-life.

### Pupil size

While our study overall implies that MIB may occur during actual self-motion, such as when driving, spontaneous reports of MIB during driving are rare. Why is that so? One possibility is that MIB phenomena are subtle so that not many people notice them, regardless of the functional significance of object disappearance. As we did not ask participants how salient their MIB experiences were, we used phasic pupil responses as a proxy for subjective salience. Phasic pupil dilation is known to increase in response to surprising, novel or unexpected stimuli^26–29^, allowing us to investigate whether there were any differences in the saliency of MIB across mask conditions.

We found that phasic pupil responses were larger in response to target disappearances than reappearances, suggesting that disappearances were more salient, consistent with previous research^26^. We further found that in the optic flow conditions, target disappearances in the static mask condition induced larger pupil dilations than both the expansion and contraction conditions. This suggests that disappearances in the static condition were more salient, which might be explained by the fact that target disappearances were much less frequent in this condition. Conversely, and importantly, this means that target disappearances are less salient during optic flow stimulation. Reappearances did not show differences in terms of pupil size between mask conditions, probably because every disappearance is predictably followed by a reappearance after a relatively short period, making them less surprising. Interestingly, the differences in pupil responses between static and moving mask conditions were not significant in Experiment 4, perhaps because the target was embedded in a mask with higher dot densities, making the disappearance less salient. Overall, the pattern of pupillary responses we observed appears consistent with an interpretation that they signal the subjective saliency or surprise of events^26–29^. Importantly for our discussion here, disappearances are less salient in optic flow conditions than those in a static scene, which may explain why people seldom notice MIB during driving.

When analysing the tonic pupil diameter (i.e. the average pupil diameter per mask condition), our results were again consistent with the phasic saliency account, though this time concerning a less-transient surprise measure that was dependent on motion mask type. Specifically, compared to the static mask condition, tonic pupil size was smaller during 3-D expanding optic flow, which is a condition that is frequently experienced during forward self-motion. Consistent with this idea, the opposite effect was found in Experiment 4, where average pupil size increased significantly for the 2-D radial expansion condition, where dots moved in an unnaturally constant speed across the screen - a situation we do not encounter in our environment.

These opposite effects of the tonic pupil diameter on the background motion pattern can be explained if we assume that they are dependent on the subjective amount of surprise. Frequently experienced 3-D expansion (and contraction) caused by self-motion are well internalized in the visual system, causing little surprise^46, 47^. The visual system does not have good internal models for unnatural 2-D expansion motion, which therefore produces greater surprise and an increase in subjective saliency.

If realistic patterns of motion are indeed less salient, and thus attract less attention, it could mean that more attention is available to be directed toward other elements in the visual display, such as the targets. If there is indeed more attention paid to the targets embedded in realistic motion types, more target disappearance should result^15, 17^. Further research on MIB is necessary to understand how surprise-related pupillary responses are modulated by the allocation of top-down attention and subjective awareness of perceptual events and by requirement of reports^48–50^. For example, one could quantify the threshold contrast increment of a target^51^. Such a probe task can be performed during a driving task without any explicit report on the target visibility itself^52^, as a function of concurrently measured pupil diameter as a proxy of subjective disappearance of the target. If indeed the threshold increases during the phasic increase of pupil dilation under the no-report condition, it has a strong implication for the existence of unreported MIB during real-life driving.

### Limitations and implications

While providing support for future studies on the occurrence of MIB in the natural world, our research was still undertaken in an artificial setting. The experiment was conducted on a computer display and some aspects of the display itself were also inconsistent with the type of motion perceived in daily life. Specifically, the mask dots did not increase in size as they approached the periphery, as would occur in normal optic flow movement^53^. Additionally, the mask was based on a tunnel, yet driving through tunnels in reality is usually brief and infrequently experienced.

Compared to previous research^1, 6^, percentages of disappearance was smaller and mean disappearance episode lengths were shorter. This is because we intentionally avoided optimising the task design to increase target disappearance. Our original research objective was to investigate if MIB could occur under naturalistic self-motion induced optic flow, and thus we did not attempt to optimise the parameters for disappearances (e.g., reducing the size or enhancing the saliency of the target for more frequent MIB). The task also had a lower density of dots around the target as a result of the expanding motion, which reduces competition between the mask and target, causing less target disappearance^1, 20^. Similarly, compared to previous studies^1^, we found a relatively high rate of Troxler fading within the static mask, a control condition for MIB. Troxler fading is known to increase with lower luminance contrast as used in our study^6^. Indeed, when compared with previous research utilising masks also not designed to optimise MIB^8^ our results were comparable or stronger in effect. Our results thus provide a conservative estimate of the potential occurrence of MIB in real-life settings.

Collectively, our findings show that optic flow simulating self-motion successfully elicits MIB. If found to occur in the real world, MIB may hold implications for driving safety, where the most common point of gaze is at the centre of the expanding optic flow of the approaching road^54–56^. Additionally, items that need to be monitored, such as brake lights of cars ahead, overtaking cars or GPS devices attached to the windscreen, reside mainly in (upper) peripheral quadrants, which we found prone to perceptual disappearance.

While drivers may often saccade to check mirrors, scan for other cars and so forth, there are also situations in which fixations lengthen and saccade frequency decreases. These include when driving on rural as opposed to urban roads^45, 57^, during times of danger or hazard^45^, when multi-tasking^58, 59^ and when experiencing fatigue^60^. Given that fewer saccades and longer fixations result in greater target disappearance, MIB may be more likely to occur during hazardous situations or driver fatigue - two of the most influential factors relating to traffic accidents^61^.

Future research should attempt to measure the occurrence of MIB directly in conditions that approximate the statistics of the real world experience in a closer fashion, especially given its potential impact on driving safety. Examples of this may be tasks which measure brightness increments and reaction time in MIB, or apply the probe technique used in Binocular Rivalry to measure MIB indirectly, where the detection of a probe is expected to be impaired during target disappearance. Tasks such as these will hold stronger implications for driving safety, where MIB may play a role in the failure to detect crucial objects or changes in appearance, such as the brake lights of a car travelling in front. Such research could utilise a virtual reality environment to directly test the hypothesis of MIB occurring under driving circumstances, which would allow for stronger conclusions of the real-world implications than can be drawn from our stimuli.

If confirmed to exist under more realistic visual settings, MIB may also hold implications for the development of Head-Up Display (HUD) technology. Using this technology crucial information is projected onto the windscreen or visor in the form of stable symbols and text, through which the moving background of road or land is viewed. Such technology can be found, for example, in the use of F-35 Fighter Jet helmets which have been developed to project flight information onto the visor viewed by the pilot over the moving background^62^, or increasingly in commercially available cars with dashboard information projected onto the windscreen^63^.

## Methods

### Participants

Ethics approval was obtained from the Monash University Human Research Ethics Committee, and informed consent was obtained from all participants in the form of written permission prior to participation. Experiments were carried out in accordance with with the relevant guidelines and regulations. Thirty-seven participants were recruited at Monash University, all meeting the requirement of normal or corrected-to-normal vision. There was no overlap between the 18 participants for Experiments 1 and 3 (aged 18–59 years, *M* = 24, *SD* = 9; 15 females) and 19 participants for Experiments 2 and 4 (aged 18–34 years, *M* = 23, *SD* = 4; 13 females).

### Apparatus

All experimental displays were created using Matlab (Mathworks, Natick, MA) and OpenGL, with the PsychToolbox extension^64, 65^. In Experiments 1 and 3, the experimental display was an IBM P275 CRT powered by a Dell Optiplex9010, and viewed from an approximate distance of 45 cm. In Experiments 2 and 4, the experimental display was a Tobii TX-300 powered by a MacBook Pro OS 10.9.5, viewed from an approximate distance of 60 cm. Experiments were run in an experimental room without natural light, with artificial lights dimmed in Experiments 1 and 3, and adjusted to a level enabling the proper performance of the Tobii TX-300 eye tracker in Experiments 2 and 4.

Hereafter, the stimulus specifications presented are for Experiments 1 and 3 with those for Experiments 2 and 4 in brackets. The stimulus consisted of a grey background at 31 Cd/m2 (31 Cd/m2), and white mask dots at 83 Cd/m2 (169 Cd/m2), each with a radius of .05° (.04°; Supplementary Movie S1). The mask was constructed as a 2-D rendering of a 3-D circular tunnel with randomly placed dots, through which the participant experienced simulated forward or backward motion. The tunnel was generated anew for each 1-minute trial, had a 5 meter radius, and was 3000 meters long. The mask consisted of 7500 white dots, placed randomly on the perimeter of the tunnel. Due to the trial length of 1 minute not all 7500 were seen by the participant. Dots moved progressively out of a central cluster (about .50° (.38°) radius), at which subjects were instructed to fixate during the experiment. During simulated forward self-motion the speed of dot movement gradually increased as each ‘moved nearer’ to the participant and approached the periphery of the screen, a pattern which was reversed for backward-motion conditions. Unlike real-life dot progression however, the dots did not change in size with expansion or contraction^53^, because we aimed to focus primarily on the influence of motion information.

The peripheral target for MIB was a white disc with a .84° (.63°) radius, surrounded by a thin exclusion boundary of .34°(.25°), through which no mask dots travelled. The disc was located 10.75° (9.32°) degrees diagonal distance from the central fixation cluster, 6.36° (4.61°) vertical and 8.70° (8.10°) horizontal, positioned in either the upper left, upper right, lower left or lower right quadrant of the screen in each trial.

Within each 60-second trial, one catch episode occurred in which the target physically disappeared from the display for a short period of time. The luminance of the target was linearly ramped off over 1.5 seconds, and stayed grey (i.e., the color of the background) for 1-4 seconds (randomly assigned in each trial), then linearly ramped up to white (i.e., the color of the original target) over another 1.5 seconds. The total duration of the catch episode therefore varied randomly from 4-7 seconds across trials. Each catch episode started randomly between 10 seconds and 50 seconds after the onset of each trial, to appear indistinguishable from perceptual disappearances. Participants were not told of the catch, but only encouraged to report all perceived disappearances. Accordingly, if the participants did not report physical disappearance of the catch appropriately, we regarded that they had not paid attention to the task in that trial. Debriefing with a number of participants revealed that they were surprised to hear of the catch disappearances, as they had not been able to tell the catch from the genuine perceptual disappearances. While three participants failed to record the catch on one or more trials, analyses with these participants excluded produced no significant difference to results. Their data was therefore included for analysis.

In Experiments 2 and 4, we also recorded participant fixation locations using an eye tracker. One purpose of eye tracking was to exclude participants who looked directly at the peripheral target, failing to fixate on the central point at any time. Fixations were defined as any duration between macro-saccades (*<*1 deg). If the fixation duration on the peripheral target per 60-second trial was longer than one second, the participant was considered for exclusion. Only three participants met this exclusion criterion on at least one trial. As including or excluding them did not significantly alter the results, we included their data for analysis. The other purpose of eye tracking was to quantify the influence of eye movements on MIB^31, 44^, which might to some degree explain the effects of our experimental manipulation of motion stimuli.

We summarise the purpose of each experiment as follows.

#### Experiments 1 and 2

In Experiments 1 and 2, three different types of mask were presented – expansion (simulated forward motion); contraction (simulated backwards motion), and a static still frame in which there was no dot movement. The simulated movement speed was 65 km/h for both expansion and contraction. The four different target locations were balanced and randomised over trials. Both Experiments 1 and 2 consisted of 12 × 1-minute trials, presenting each possible combination of mask (3 types) and target location (4 quadrants). Eye-movements were recorded during Experiment 2.

#### Experiment 3

In Experiment 3, only forward motion was simulated, resulting in an expanding motion pattern. Four different movement speeds were simulated: 35 km/h, 50 km/h, 65 km/h and 80 km/h. Experiment 3 consisted of a total of 16 × 1-minute trials, presenting each possible combination of speed and target location.

#### Experiment 4

Experiment 4 had two purposes: 1) to compare coherent radial expansion/contraction motion conditions to a condition with 2D random motion and 2) to investigate whether MIB still occurred as strongly when the mask dots moved at a constant speed on a 2D plane, unlike those that moved in an accelerating speed as in the optic flow conditions in 3D space (as in Experiment 1-3) In Experiment 4, instead of a central cluster, the fixation point was explicitly presented as a white cross with a size of 0.50° x 0.50° (Supplementary Video S2). The cross was surrounded by an exclusion radius of 2.35°, through which no mask dots travelled. In addition, only in Experiment 4, all dots (500 in total) moved at a constant speed of 3.4° per second. Four different types of mask were presented: 2-D radial expansion, 2-D radial contraction, 2-D static and 2-D random movement. With the random type of mask movement, the dots moved in a random direction and had a 1% chance of being removed on each frame. Removed dots, and dots which crossed the outer edge of the display were replaced at a random position on the screen. Experiment 4 consisted of 16 × 1-minute trials, presenting each possible combination of mask type and target location. Due to the stimulus layout, this experiment had a uniform density of mask dots on the screen, while the other experiments had a decreasing density with increasing eccentricity.

### Procedure

#### Experiments 1 and 3

Participants sat comfortably in front of the computer, with their head stabilised by a chin rest. They were instructed to fixate on the central area of the screen and to indicate the perceptual invisibility of the target by holding down the spacebar for as long as the target remained invisible. Participants were instructed to pay attention to the peripheral target, without looking at it directly.

#### Experiments 2 and 4

Experiments 2 and 4 followed the same procedure as 1 and 3, however eye movements were also recorded. Participants underwent a standard 9-point eye-tracking calibration before the experiment started, and did not move their heads away from the chin rest for the duration of the experiment.

### Analyses

We excluded the catch events from our analysis for the calculation of percentage disappearance, rate, and durations of disappearance episodes. For percentage disappearance we excluded the part of reported disappearances that overlapped with the catch, and total trial duration was adjusted to exclude the catch event. For rate, we excluded disappearances that started during the catch event. For the durations of disappearance episodes we excluded disappearances that overlapped the catch by any amount, and we also excluded disappearances that overlapped with the end of the trial.

We ran three one-way repeated measures ANOVAs with mask condition as the sole independent variable (IV) for each experiment. The mean percentage, rate or duration of disappearance per condition were the three dependent variables (DV). For comparison, all analyses were also run using the median instead of the mean. The results did not change in a significant way (see Table 1).

To examine how much of the observed MIB disappearance (in each trial) could be explained by the frequency of saccades, which are expected to reduce MIB, and the location of the target, we used the linear mixed effect model (LME) in Matlab (ver 2016a). Likelihood ratio tests were performed through a c^2^ test comparing the full model and a reduced model without the factor or interaction in question^32^. We regarded percentage, rate or duration of disappearance as DV and modeled DV in each trial with a fixed effect of MaskType (for the different types of motion mask), NumberOfSaccades (per trial), and the target location (LeftRight and UpDown) as DV MaskType + NumberOfSaccades + LeftRight + UpDown + (1| SubjectID). Subject variability was taken into account as a random effect. NumberOfSaccades was only entered when available (Experiment 2 and 4), in which case we included it in the unrestricted model as an interaction with MaskType.

Assumptions for all tests were met, with the exception of violated normality in several conditions across experiments. Unless otherwise stated a square root data transformation was performed for all ANOVA analyses, as small sample size (*<*30) meant the Central Limit Theorem was not applicable in correcting normality^66–68^. A constant of 1 was added to all data when performing the transformation, due to the dataset containing 0 values^66^.

Despite the transformation, normality was occasionally still violated in the static mask condition across Experiments 1, 2 and 4, with skewness Z-Scores exceeding the alpha =.05 cutoff of 1.96^66, 67^. Alternate transformations (Log10, Reciprocal) also failed to correct normality in this variable. The violation in the static mask control condition was due to the majority of participants recording disappearance scores of zero for this condition, as expected. Given that negative values were not possible in our study, this necessarily led to a skewed distribution in the static condition. As the violated static condition was serving as a control for comparison only within our ANOVA analyses, the square root transformation was retained and parametric tests run despite the violation. However, non-parametric Friedman’s tests corroborated our findings in all cases. All graphical representations of data were created using within-subjects error bars displaying +/- 1 SE, as recommended by Cousineau^69^.

### Eye-movement analyses

#### Saccades and fixations

Eye movements were recorded binocularly using a Tobii TX-300 Eyetracker (Tobii Technology, Danderyd, Sweden) at a sampling rate of 300Hz, controlled through the software package T2T (http://psy.cns.sissa.it/t2t/AboutT2T.html). Saccades and fixations were detected offline using a velocity-based algorithm^30^. Saccades with an amplitude larger or smaller than 1 degree of visual angle were categorised as macro-saccades or micro-saccades, respectively.

#### Pupil size

Pupil diameter was also recorded by the Tobii TX-300 Eyetracker. We analysed the left eye and followed procedures by Kloosterman et al^26^. Briefly, to obtain phasic pupillary responses, we took excerpts from the pupil trace from −1 second to + 1.5 seconds around the time of a reported disappearance or reappearance. The pupil diameter over the first 400 ms of each excerpt was subtracted from the signal, and the result was then divided by the mean pupil diameter over the entire experiment (dividing by the mean per condition yielded identical results in terms of passing statistical significance). This was done per participant separately. Results were then analysed over participants, with multiple comparisons corrected with a False Discovery Rate (FDR) 0.05. To estimate the sustained, baseline tonic level of pupillary diameter for each motion stimulus, we calculated the mean pupil diameter for each of the movement conditions separately, without normalisation.

## Author contributions statement

VT and JVB conceptualised and designed the research reported in this article, with feedback from NT and MD. VT collected and analysed the behavioural data, with help from JVB. VT wrote the article, with feedback from JVB, MD and NT. MD programmed the experiments, with help from JVB. PZ analysed the saccade data, JVB analysed the pupil data.

